# Braintree: An agentic framework for in-silico vision neuroscience

**DOI:** 10.64898/2026.10.07.756448

**Authors:** Mainak Deb, Ruolin Wang, Alish Dipani, N. Apurva Ratan Murty

## Abstract

Vision neuroscience has many of the ingredients needed to conduct experiments entirely in-silico. We need AI systems that can reason over scientific knowledge, generative methods for creating precisely controlled stimuli, image-computable models that can predict neural responses with increasing accuracy, and tools that can analyze and interpret data. Here we take the first step toward making a vision neuroscience experiment end-to-end computable. We introduce Braintree, an agentic framework that can autonomously run multi-experiment studies directly with in-silico models of human visual cortex. Given a hypothesis, Braintree can reason over the current literature, design an experiment, build a stimulus set from algorithmically manipulated or synthesized stimuli, run the experiment on candidate brain encoding models, analyze the responses and then use the results to design even stronger tests. The entire loop runs autonomously and can accept human input at any stage. We evaluated Braintree on three increasingly open-ended scientific problems. First, Braintree reproduced published findings using their original stimuli and then asked whether the same effects could be replicated with entirely new stimulus sets that it constructed. Second, it designed new experiments to find stimulus dimensions that could drive two brain regions differently, even when those regions otherwise had similar response profiles. Third, it searched for experiments that could tell competing models of the same brain region apart. Experiments discovered using just two models also exposed differences across a much broader set of models, suggesting that Braintree had found more general dimensions along which brain models disagree. Braintree can augment the capabilities of a vision neuroscientist as a verifier for existing hypotheses, a test-bed for new ideas, and/or a tool for generating stimuli that can discriminate between competing hypotheses or models of the brain. Braintree shows how AI agents and brain models can work together to make in-silico vision neuroscience experiment computable, allowing vision scientists to explore many more experiment conditions and identify the most informative ones to test on the human brain.

## 1 Introduction and Related Work

Vision neuroscience is on the cusp of becoming *experiment-computable*. Given a specified study, we can now execute it directly on in-silico models of the human brain (Yamins & DiCarlo, 2016; Schrimpf et al., 2020; Conwell et al., 2024; Gifford et al., 2025a; Wang et al., 2026). We can present arbitrary images, measure predicted neural responses, and repeat this process across thousands or even millions of experimental conditions (Ratan Murty et al., 2021; Gifford et al., 2025b). We also have powerful methods for generating and manipulating visual stimuli, which give vision scientists precise control over stimulus features and the ability to isolate variables and rule out confounds (Walker et al., 2019; Ponce et al., 2019; Luo et al., 2023; Garcia Cerdas et al., 2025). But we do not have an existing framework that can make the *experiment design process* (ideation, planning, stimulus synthesis, controls, data analysis, etc.) scale with our newfound (in-silico) *experiment capacity*.

Other areas of science are beginning to use agentic systems to scale experimentation and discovery (for example, see (Lu et al., 2024; Ghareeb et al., 2026; Gottweis et al., 2025)). These systems can help formulate hypotheses, design and execute computational studies, analyze their outcomes, and in some cases, use the results to guide subsequent experiments. Agentic approaches are emerging (but are still nascent) in neuroscience. Some recent agentic systems support specific stages of the scientific pipeline like hypothesis generation, data preprocessing, analysis, and interpretation (Li et al., 2026a; Wu et al., 2026; Li et al., 2026b). Vision neuroscience is also unique in the sense that both the study design and study execution sides could be made autonomous. We have the ability now to construct and precisely manipulate images that can go into an experiment, and image-computable in-silico brain models can predict the neural responses to those images. It is therefore in principle possible to turn a scientific question into an experiment, generate the stimuli rule out confounds, measure the resulting brain responses, and use those responses to determine what experiment should come next. Can we bring these capabilities together into a closed experimental loop?

In this work, we introduce Braintree, an agentic framework for carrying out the full cycle of a vision neuroscience experiment, from scientific question to experiment planning, stimulus design, study execution, data analysis and inference. A scientist begins with a research question and provides as much or as little guidance as needed. Braintree turns this input into an experimental plan, designs and verifies the stimuli, runs the experiment on models of the visual brain, analyzes the resulting responses, and uses those results to guide what to test next. This process is carried out by specialized agents for planning, stimulus design, experimentation, analysis, and scientific inference, with dedicated auditing agents that verify each stage. The scientist can inspect, modify, or redirect the study at any point. Braintree therefore provides a way to move from a scientific question to a sequence of executable experiments, while keeping the scientist in control throughout.

We describe the structure of the Braintree pipeline and then use it to demonstrate three capabilities. *First*, we show that the full pipeline can reproduce and then replicate findings across dozens of published vision neuroscience studies without human intervention. *Second*, we show that Braintree can go beyond existing studies and design new experiments to find stimulus manipulations that dissociate pairs of brain regions with otherwise similar response profiles. *Third*, we show that Braintree can design experiments that expose disagreements between brain models, including experiments discovered using two models that generalize to a broader population of models. Together, these results show how agentic systems can turn in-silico brain models from passive predictors of neural responses into systems that can be actively experimented on.

## Methods

### Braintree framework design

A vision neuroscience study naturally decomposes into four stages, which we formalize as protocols (Fig. 1). The *Knowledge* protocol translates a scientific question into a grounded experimental goal. The *Design* protocol turns this goal into an executable experiment, including the design and stimuli. The *Execution* protocol runs the experiment on in-silico models of the human visual cortex and predicts neural responses. Finally, the *Inference* protocol analyzes these responses and draws a scientific inference. We formalize this cycle as

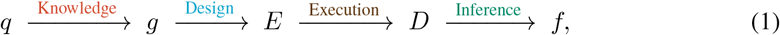

where *q* is the scientific question, *g* is the grounded experimental goal, *E* is the executable experiment, *D* is the resulting predicted neural response data, and *f* is the scientific inference. The inference *f* either concludes the study with a verdict or seeds the next cycle as a refined question *q*′.

**Figure 1:**
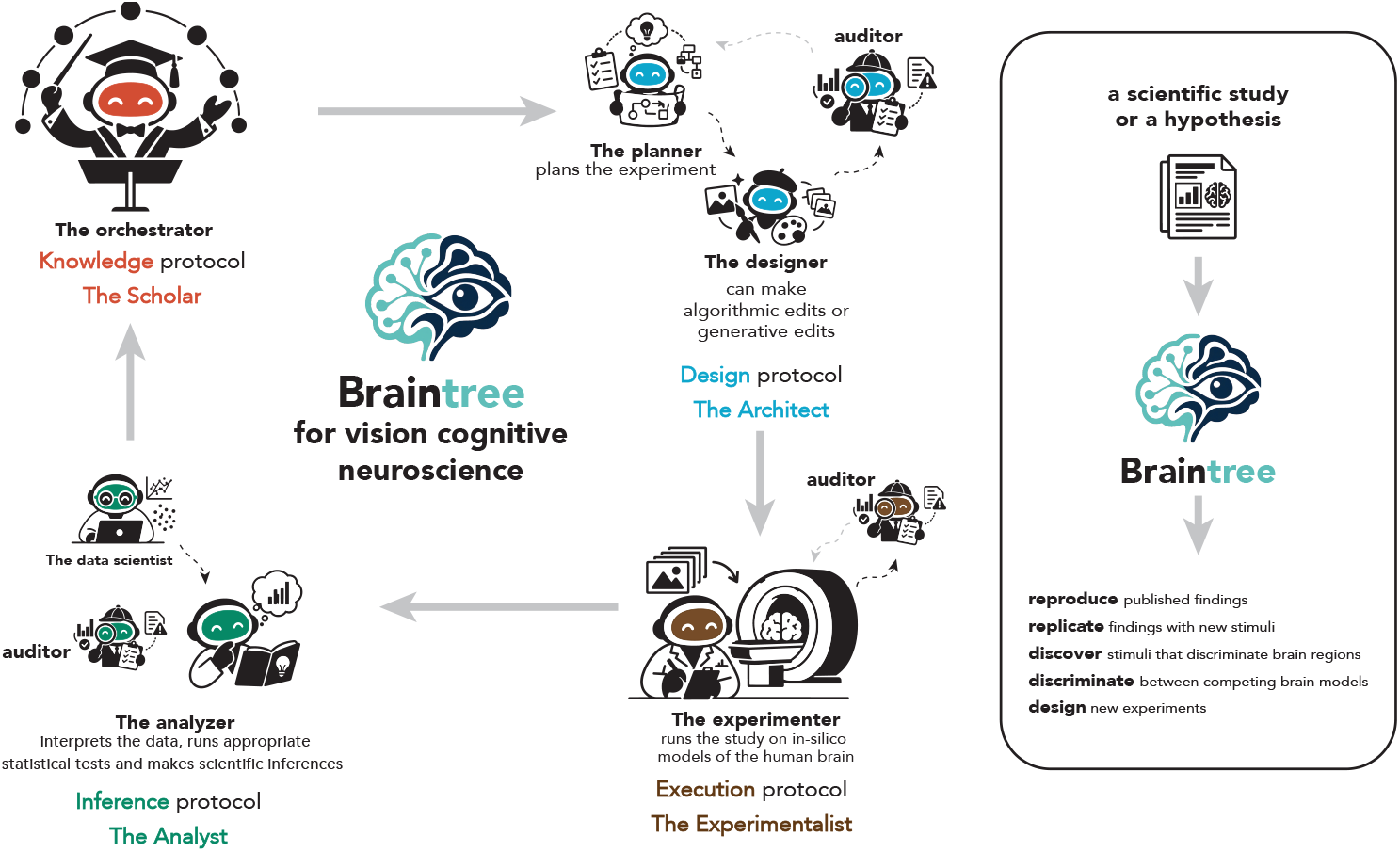
Braintree: an agentic framework for vision neuroscience. Braintree organizes the experimental process into four protocols: Knowledge, Design, Execution, and Inference. A central orchestrator coordinates specialized agents that plan experiments, construct stimuli using algorithmic or generative edits, execute studies on in-silico models of the human brain, and analyze the resulting responses to draw scientific inferences. Independent auditor agents verify intermediate outputs throughout the pipeline. Braintree can reproduce published findings, replicate them with new stimuli, discover stimuli that discriminate between brain regions or competing brain models, and design new experiments.

Braintree implements this cycle with a set of specialized agents (Fig. 1; Table 1). An orchestrator agent coordinates them and serves as the user’s interface to the system, so the process can be steered at any point. A planner agent and a designer agent produce the experimental design and construct the stimuli; an experimenter agent runs the experiment on image-computable in-silico models of the human brain, and an analyzer agent interprets the predicted responses and draws inferences that guide subsequent experiments. Independent auditor agents verify the experimental design, stimuli, results, and inferences, and all quantitative analyses are performed by fixed analysis code rather than by language models. Our current implementation instantiates the agents with the Claude family of models (see Table 2 for the full list). Further details of the agents, models, statistical procedures, and auditing are given in Appendix A.1.

**Table 1:** Main agent roles used by Braintree. Agents have narrow responsibilities so that experimental design, execution, and evaluation can be checked independently.

| Agent role | Responsibilities |
| --- | --- |
| Orchestrator | Maintains the scientific goal and coordinates the experimental study. |
| Planner | Converts the scientific question into an executable experimental design. |
| Designer | Constructs or selects stimuli implementing the experimental conditions. |
| Auditor | Independently checks stimuli and experimental specifications before execution. |
| Experimenter | Runs validated stimuli through the in-silico brain models. |
| Analyzer | Summarizes and visualizes the resulting neural responses. |

**Table 2:** Models behind the agents of Figure 1. All agents are Claude Code subagents on Anthropic’s Claude 5 family; sampling parameters are the API defaults with adaptive thinking on, and the effort level is the only depth control. The orchestrator of the region-dissociation experiments (Section 3.2) ran on Claude Opus 5 at high effort. The data scientist is deterministic analysis code and calls no language model.

| Protocol | Agent | Model tier | Effort |
| --- | --- | --- | --- |
| Knowledge | Orchestrator | Claude Opus 5 | high |
| Design | Planner | Claude Sonnet 5 | low |
| Design | Designer | Claude Sonnet 5 | low |
| Design | Auditor | Claude Opus 5 | medium |
| Execution | Experimenter | Claude Haiku 4.5 | low |
| Execution | Auditor | Claude Opus 5 | medium |
| Inference | Data scientist | fixed analysis code | – |
| Inference | Analyzer | Claude Sonnet 5 | low |
| Inference | Auditor | Claude Opus 5 | medium |

#### Stimulus design

Braintree supports stimulus design using (1) stimulus retrieval and curation, (2) *algorithmic* edits and (3) *generative* edits. **Stimulus retrieval and curation** searches existing image sets and finds stimuli that satisfy the constraints of an experiment. In our current implementation, the designer agent curated more than 320, 000 stimuli from 28 image datasets. **Algorithmic edits** apply deterministic or parametrically controlled manipulations to existing images. These edits cover, for example, image cropping, masking, spectral filtering, compositing, or manipulating specific image statistics. **Generative edits** use generative models to synthesize new stimuli or manipulate existing images, such as adding, removing, or modifying objects and attributes. For each experiment, the Designer selects and combines these tools, and can retrieve and integrate additional tools from the web when the required manipulation falls outside the current set.

#### In-silico brain models

Our current implementation operates on whole-brain image-computable encoding models fit to the Natural Scenes Dataset (NSD) (Allen et al., 2022). For each NSD participant *s*, the encoder implements a mapping

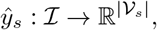

where *I* is the space of RGB images and *V*_*s*_ is the set of modeled cortical voxels for participant *s*. Given an image *x*, a pretrained vision model *ϕ* first maps the image to a feature representation *ϕ* (*x*). In most cases (unless specified) we use the CLIP-ResNet50 (Radford et al., 2021) or DINOv2 (Oquab et al., 2023) encoders as our feature models. Both are popular choices for modeling visual representations. A learned brain-mapping function *g*_*s*_ then maps these features to the predicted voxel responses,

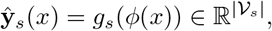

with *ŷ*_*s,v*_(*x*) denoting the predicted response of voxel *v*. The mapping *g*_*s*_ is fit once using NSD image–response pairs and is subsequently frozen. Thus, every image generated or selected by Braintree can be converted directly into a predicted whole-brain response without experiment-specific fitting.

For experiments targeting a region of interest *R*, we obtain the predicted regional response by averaging over the corresponding voxel responses,

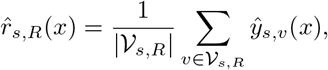

where *V*_*s,R*_ ⊆ *V*_*s*_ is the set of voxels assigned to region *R* in participant *s*. This provides Braintree with a common experimental interface: an experiment specifies a set of images 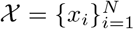and a target region *R*, and the encoder returns the predicted response vector

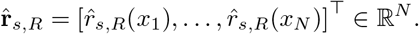

Our current implementation focuses on category-selective regions in the higher-level visual cortex (FFA, PPA, EBA, OFA, and RSC). We primarily operate at the region level because hypotheses in vision neuroscience are typically formulated and tested around specific brain regions (very rarely on the whole brain). Further details about the in-silico models and fMRI data are provided in Appendix A.2.

## 3 Results

### 3.1 Braintree can reproduce and replicate published findings

Do different parts of the Braintree pipeline work together to recover established findings from vision neuroscience? We showcase the ability of Braintree in two stages. We start with a paper **reproduction** test and then move to a paper replication test. The Paper reproduction test asks whether, given a published study and the original stimuli, Braintree can infer the goal of the experiment, execute it on an in-silico brain model, and recover the reported findings.

#### Braintree pipeline

For the paper reproduction tests, the orchestrator agent reads the original study and extracts the claim and relevant experimental details. The planner agent converts these into an executable experiment. Because the original stimuli are available, the designer agent has a limited role and only organizes the original stimuli into the appropriate experimental conditions. After the design and stimuli are audited, the experimenter agent runs the study on the corresponding in-silico brain model. The resulting responses are passed to the analyzer agent, which invokes the fixed analysis pipeline and interprets the results in the context of the original claim.

#### Results

We looked for published studies for which the original experimental stimuli were available in the public domain. We identified 10 such studies (Appendix A.3, Table 3). The input to Braintree was deliberately sparse. It was given the paper and only asked to reproduce the experiment using the available original stimuli. From there, Braintree had to determine what finding was being tested, reconstruct the relevant experimental design and conditions, organize the stimuli, run the experiment on the appropriate in-silico brain model, and analyze the resulting responses. Braintree reproduced the reported finding in *all 10 studies* (Fig. 2).

**Table 3:**
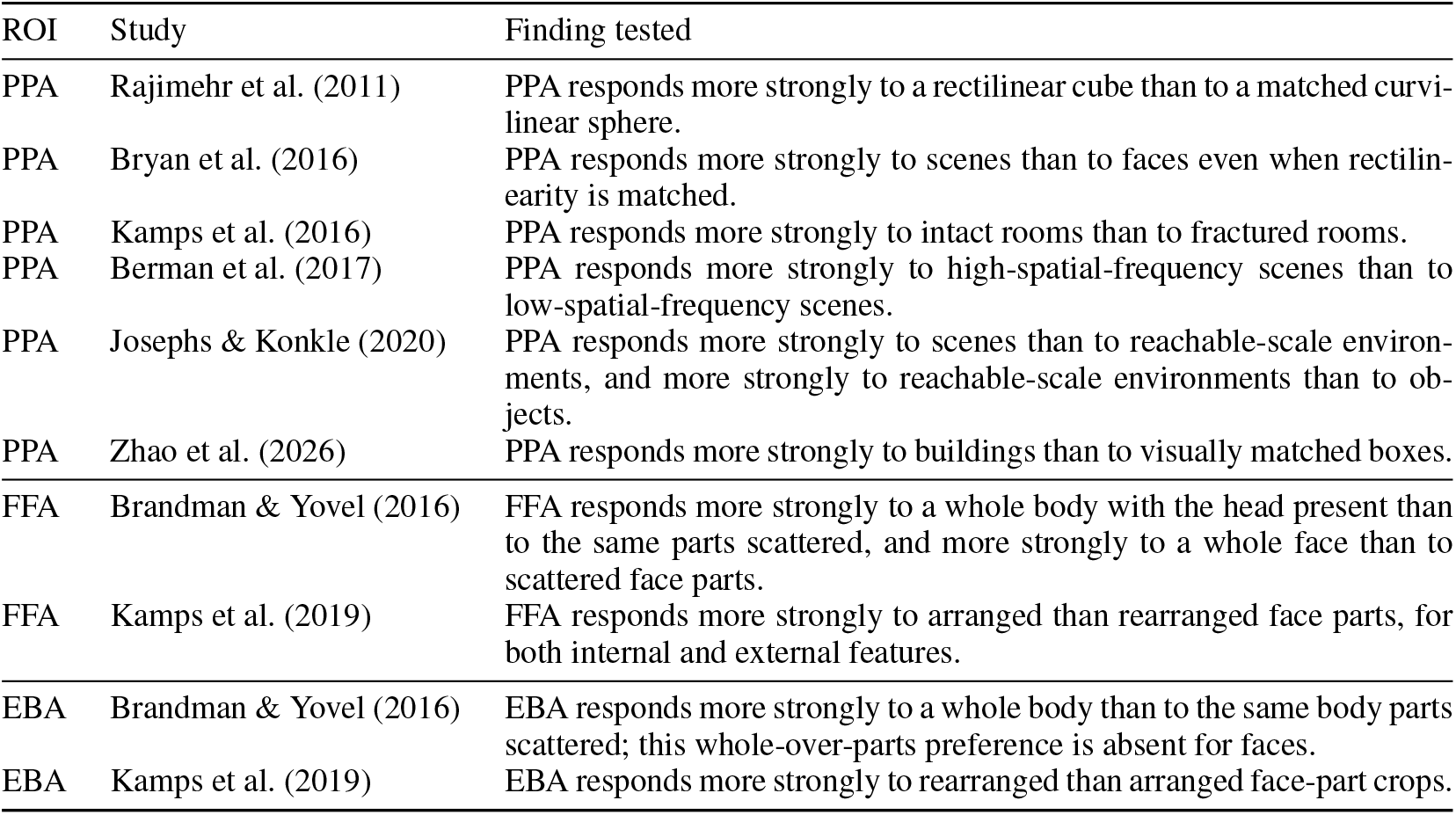
Published findings reproduced by Braintree. For each study, we report the target brain region and the principal experimental finding tested by Braintree on the authors’ original stimuli.

**Figure 2:**
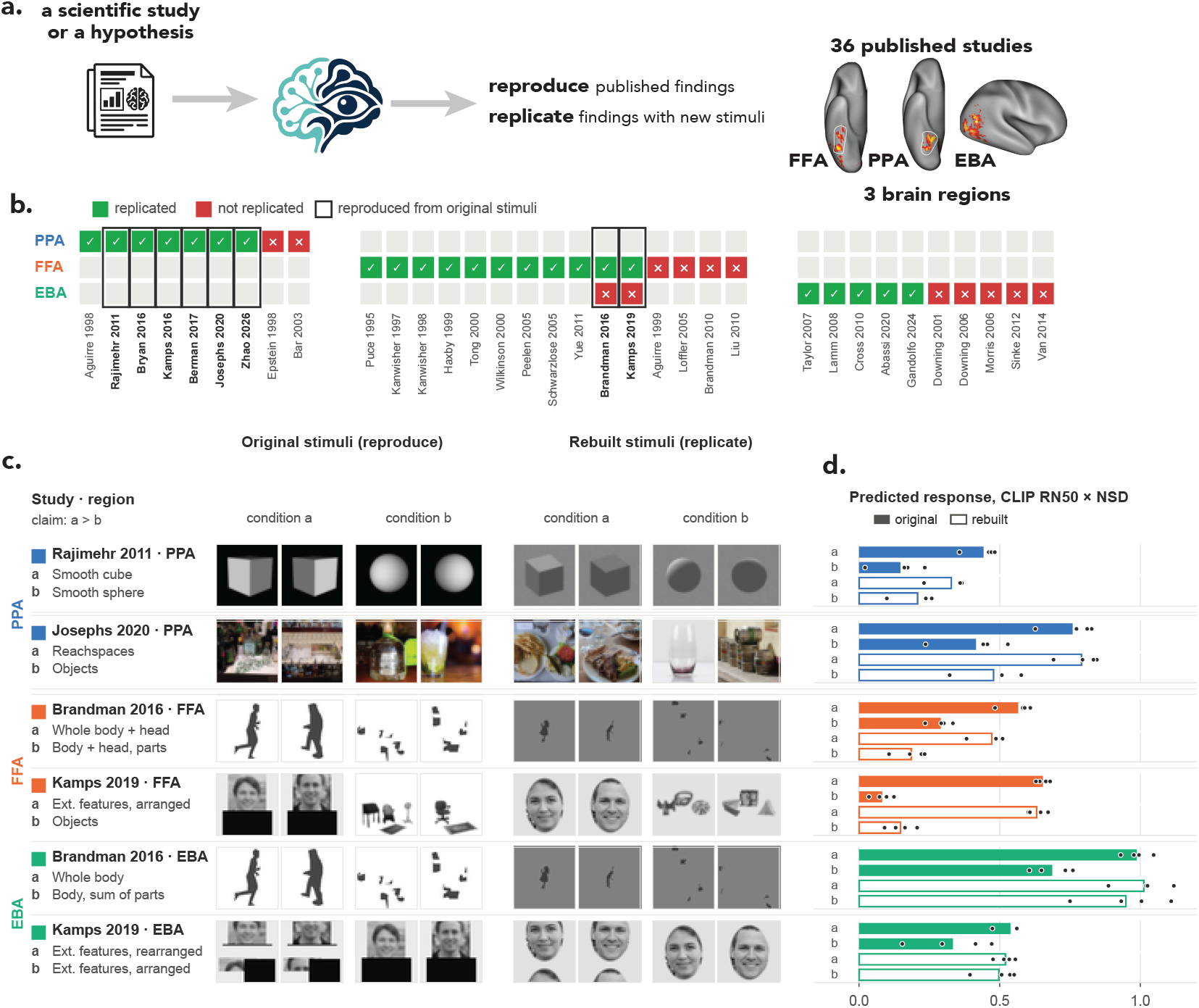
Braintree reproduced and replicated published findings in vision neuroscience. **a.** We tested 34 published findings across 3 brain regions. Reproduction used the original stimuli; replication required Braintree to construct new stimuli testing the same claim. **b**. Results across studies. Green/red indicate successful/failed replication. Black outlines mark studies with available original stimuli; all reproduction tests were successful. **c**. Example studies showing the scientific comparison, original stimuli, and new stimuli constructed by Braintree. Faces shown in this figure are synthetically generated (Hafezi, 2024) and bodies are shown as silhouettes. **d**. Predicted responses from CLIP ResNet50-based encoding models fit to NSD. Filled and outlined bars show responses to original and rebuilt stimuli, respectively; points show individual model readouts. Braintree recovered the relevant response patterns with both stimulus sets.

This simple proof-of-principle test gives us some confidence that different parts of the Braintree pipeline work together as intended. The framework can take a relatively open-ended instruction and turn it into an executable experiment and recover the published result. But of course, reproducing the experiment is a relatively easy test. The stimuli provided already suggest the choices that define the experiment. So we pushed Braintree further. We removed the original stimuli and asked whether it could *replicate* published findings by constructing a new experiment using new stimuli that test the same scientific claim.

#### Braintree pipeline

For a paper replication test, Braintree must use the logic of the original experiment but build an independent stimulus set from scratch. The orchestrator agent again extracts the claim and experimental constraints, but the planner agent must now determine what images would constitute a valid new test. Depending on the study, the designer agent retrieves and curates new images, makes controlled algorithmic manipulations, or generates new stimuli. An independent auditor checks that the resulting stimulus set implements the intended comparison before the experimenter agent runs the experiment. The resulting responses are passed through the fixed analysis pipeline and interpreted by the analyzer agent as before.

#### Results

We next asked whether Braintree could replicate the same findings without access to the original stimuli. We started with the 10 studies that Braintree had successfully reproduced, removed the original stimulus sets, and asked Braintree to construct new stimuli that independently tested each published finding. Braintree was able to successfully replicate 8/10 studies. Interestingly, the effect sizes were generally smaller for the new stimuli compared to the original images used in the studies. This pattern was true in 7/10 studies (mean reproduced 0.32 su vs. replicated 0.28 su; mean difference Δ = 0.04 su) but the difference did not reach significance with these 10 tests (one-sided Wilcoxon signed-rank test, *W* = 40, *p* = 0.12; bootstrap 95% CI [−0.03, 0.12]). We then asked how far we could scale this replication test. So we expanded the analysis to 26 additional published studies on the PPA, FFA, and EBA for which the original stimuli were not publicly available. For each study, Braintree constructed a new stimulus set and tested whether the published effect could be validated on entirely new stimuli. 23/36 (64%) effects replicated (Fig. 2; Appendix A.3, Tab. 4) without human intervention. This result points to the *first* broader use for Braintree: asking how robust the effects in the original papers are to the specific stimuli used in the study.

**Table 4:** Published findings replicated by Braintree. For each study, we report the target brain region and the principal experimental finding tested by Braintree.

| ROI | Study | Finding tested |
| --- | --- | --- |
| PPA | Aguirre et al. (1998) | PPA responds more strongly to buildings than to faces or objects. |
| PPA | Rajimehr et al. (2011) | PPA responds more strongly to a rectilinear cube than to a matched curvilinear sphere. |
| PPA | Bryan et al. (2016) | PPA responds more strongly to scenes than to faces even when rectilinearity is matched. |
| PPA | Kamps et al. (2016) | PPA responds more strongly to intact rooms than to fractured rooms. |
| PPA | Berman et al. (2017) | PPA responds more strongly to high-spatial-frequency scenes than to low-spatial-frequency scenes. |
| PPA | Josephs & Konkle (2020) | PPA responds more strongly to scenes than to reachable-scale environments, and more strongly to reachable-scale environments than to objects. |
| PPA | Zhao et al. (2026) | PPA responds more strongly to buildings than to visually matched boxes. |
| FFA | Puce et al. (1995) | FFA responds more strongly to intact faces than to luminance-matched scrambled faces. |
| FFA | Kanwisher et al. (1997) | FFA responds more strongly to faces than to objects or houses and more strongly to intact than scrambled faces. |
| FFA | Kanwisher et al. (1998) | FFA responds more strongly to upright than inverted two-tone Mooney faces. |
| FFA | Haxby et al. (1999) | Face inversion produces only a small change in the response of face-selective fusiform cortex. |
| FFA | Tong et al. (2000) | FFA responds strongly across different face types, including human and cat faces, relative to objects. |
| FFA | Wilkinson et al. (2000) | FFA responds more strongly to concentric gratings than to radial or parallel gratings. |
| FFA | Peelen & Downing (2005) | Face-selective fusiform cortex responds strongly to headless human bodies as well as faces. |
| FFA | Schwarzlose et al. (2005) | FFA shows stronger selectivity for faces than for bodies, body parts, cars, or objects. |
| FFA | Yue et al. (2011) | FFA responses to faces vary systematically with low-level properties including stimulus size and contrast. |
| FFA | Brandman & Yovel (2016) | FFA responds more strongly to a whole body with the head present than to the same parts scattered, and more strongly to a whole face than to scattered face parts. |
| FFA | Kamps et al. (2019) | FFA responds more strongly to arranged than rearranged face parts, for both internal and external features. |
| EBA | Taylor et al. (2007) | EBA responses become increasingly body-selective as progressively more of the body is shown. |
| EBA | Lamm & Decety (2008) | EBA responds similarly to images of hands undergoing painful and matched non-painful events. |
| EBA | Cross et al. (2010) | EBA responds more strongly to contorted body postures than to ordinary body postures. |
| EBA | Abassi & Papeo (2020) | EBA responds more strongly to two bodies facing each other than to the same bodies facing away. |
| EBA | Gandolfo et al. (2024) | EBA responds more strongly to facing than non-facing human body dyads. |

These results show that the full Braintree pipeline can operate end-to-end without human intervention. The replication tests especially show that the individual components of Braintree can work together to recover the scientific question, design and construct the experiment, run it on an in-silico brain model, analyze the responses, and evaluate the result. In a single day, Braintree was able to reproduce and replicate findings across dozens of published vision neuroscience studies.

### 3.2 Braintree finds stimulus manipulations that dissociate brain region pairs

Next we asked whether Braintree can go beyond the existing literature and design new experiments to answer some open scientific questions. We showcase this capability by asking Braintree to find stimulus pairs that strongly dissociate pairs of brain regions that otherwise appear to respond to similar visual stimuli. For example, the FFA and OFA. Both regions respond strongly to faces. Similarly, the PPA and RSC both respond selectively to scenes. What kind of stimuli might drive one region more than the other. Answering this experimentally is hard because there is a very large space of stimulus dimensions one could test.

Braintree pipeline: Here Braintree was given a pair of brain regions and asked to find an interpretable stimulus manipulations that dissociates their responses. The orchestrator agent identifies what is known about the two regions and uses this information to propose candidate dimensions along which their responses might differ. The planner agent turns a candidate into an experiment, and the designer agent constructs images that vary the proposed dimension while controlling other factors. After the stimuli were independently audited, the experimenter agent runs the experiment on both regions. The resulting responses were passed through the fixed analysis pipeline and interpreted by the Inference protocol. Braintree then used the results to refine its hypothesis and design the next experiment, repeating this process until it found multiple manipulations.

#### Results

Braintree found many ways to dissociate both pairs of brain regions (Fig. 3). For FFA and OFA, it tested 28 manipulations, of which 11 produced a statistically significant dissociation (region × condition interaction beyond 0.10 standard units in every subject; *t*-tests, all *p <* 10^−11^; Fig. 3a; Appendix A.4). Interestingly, Braintree naturally focused much of its search on faces, testing different ways of manipulating face information. The dissociations also went in both directions. Low-pass filtering of faces, for example, preferentially changed FFA responses (*D* = +0.22, *p* = 1.3 × 10^−12^), whereas high-pass filtering preferentially changed OFA responses (*D* = −0.21, *p* = 1.0 × 10^−39^; Fig. 3b).

**Figure 3:**
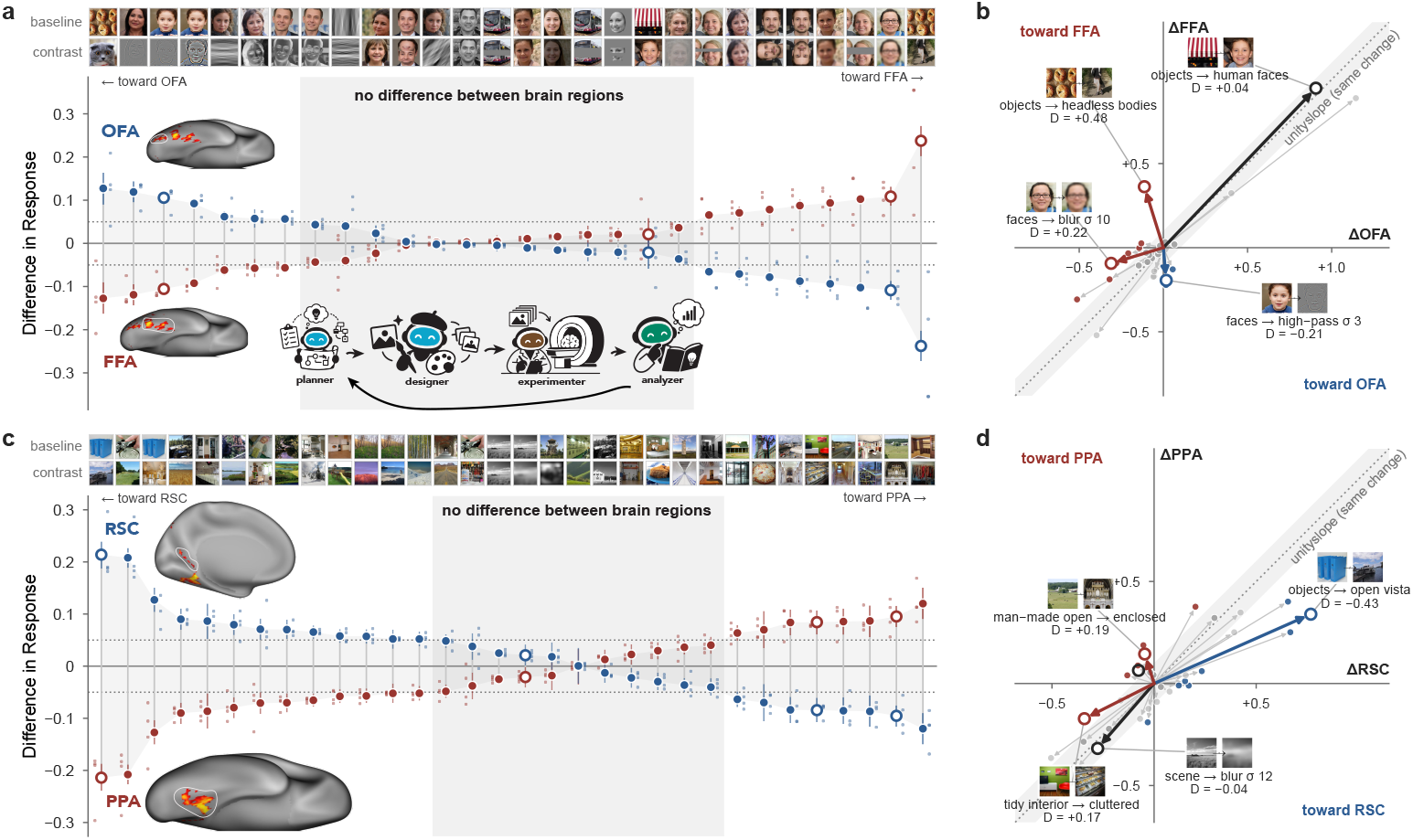
Braintree finds stimulus manipulations that dissociate brain regions. **a**. Braintree searches for interpretable stimulus manipulations that differentially modulate responses in FFA and OFA. Each point shows the difference in predicted response between a baseline and contrast condition. Manipulations are ordered by the degree and direction of the resulting dissociation, from those favoring OFA (left) to those favoring FFA (right). Example baseline and contrast stimuli are shown above. Human faces shown in this figure are synthetically generated. The schematic illustrates the iterative experimental loop in which Braintree designs, executes, and analyzes experiments and uses the results to guide subsequent experiments. Open circles indicate selected manipulations highlighted in panel b. **b**. Each manipulation is plotted by its effect on OFA (ΔOFA) and FFA (ΔFFA), revealing manipulations that preferentially affect one region. Example manipulations producing strong dissociations are shown. The diagonal indicates similar effects across the two regions. **c**. Same as panel a, for PPA and RSC, with manipulations ordered from those favoring RSC (left) to those favoring PPA (right). **d**. Same as panel b, showing the effects of individual manipulations on RSC (ΔRSC) and PPA (ΔPPA).

**Figure 4:**
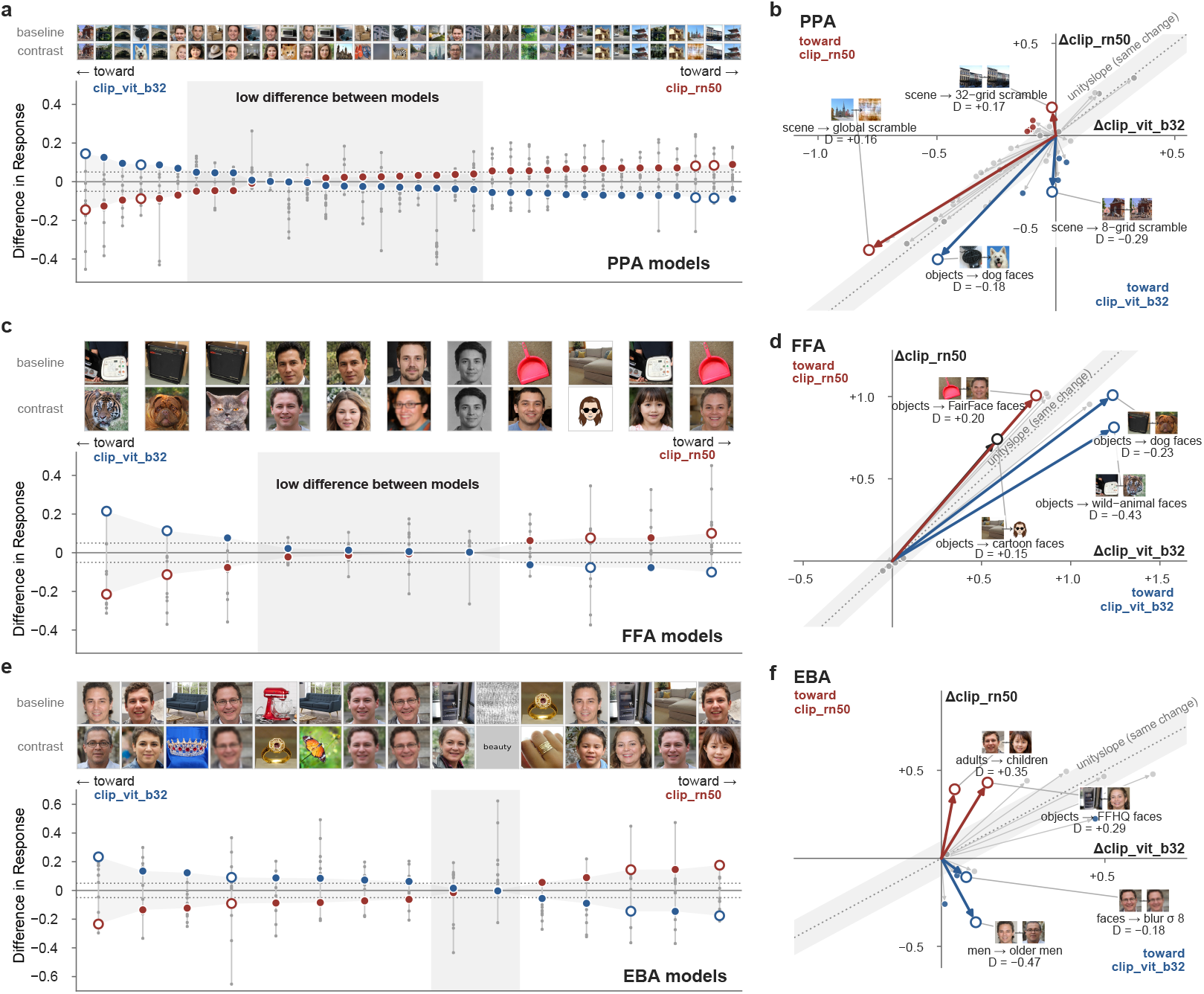
Braintree finds experiments that distinguish competing brain models. **a,c,e**. Braintree searched for stimulus manipulations that dissociated CLIP-RN50 and CLIP-ViTB32 encoding models of PPA (**a**), FFA (**c**), and EBA (**e**). Each manipulation shows the difference in predicted response between the two models, ordered from those favoring CLIP-ViT-B32 (left) to CLIP-RN50 (right). Gray regions indicate experiments for which the models responded similarly. Example baseline and contrast stimuli are shown above. Human faces shown in this figure are synthetically generated. **b**,**d**,**f**. Responses of the two models to selected experiments for PPA (b), FFA (**d**), and EBA (**f**). Each line shows the change in predicted response from baseline to contrast for the two models; the unity line indicates identical changes. Braintree found interpretable experiments that produced disagreements in both directions, such as spatial scrambling in PPA, different face categories in FFA, and age in EBA.

The hypotheses tested looked very different for the PPA and RSC pair. Here, Braintree explored properties related to natural scenes like openness, enclosure, and indoor versus outdoor setting rather than low-level image manipulations (Fig. 3c). Of the 32 manipulations tested, 7 significantly dissociated the regions (same criterion as before; paired *t*-tests, all *p <* 10^−10^). Again, the effects went in both directions. Enclosed vs open man-made scenes produced one of the strongest PPA-favoring effects (*D* = +0.19 su, *p* = 8.3 × 10^−17^), while open vistas vs objects produced a strong RSC-favoring effect (*D* = −0.43 su, *p* = 1.1 × 10^−30^; Fig. 3d). A detailed preview of the stimuli is provided in Appendix A.4, Fig. 5 and Fig. 6.

**Figure 5.**
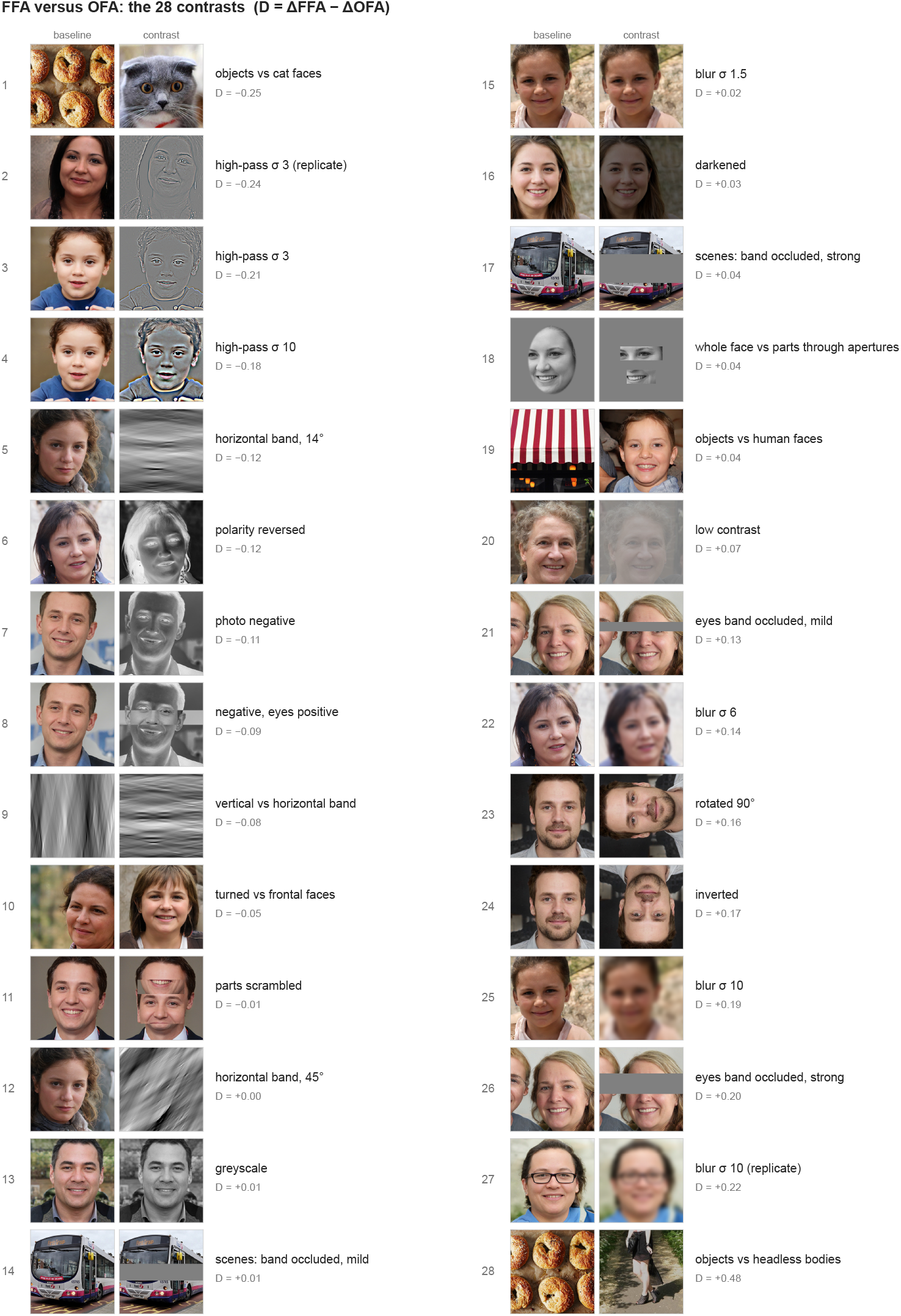
Braintree discovers stimulus manipulations that dissociate brain regions FFA and OFA. Human faces shown in this figure are synthetically generated.

**Figure 6.**
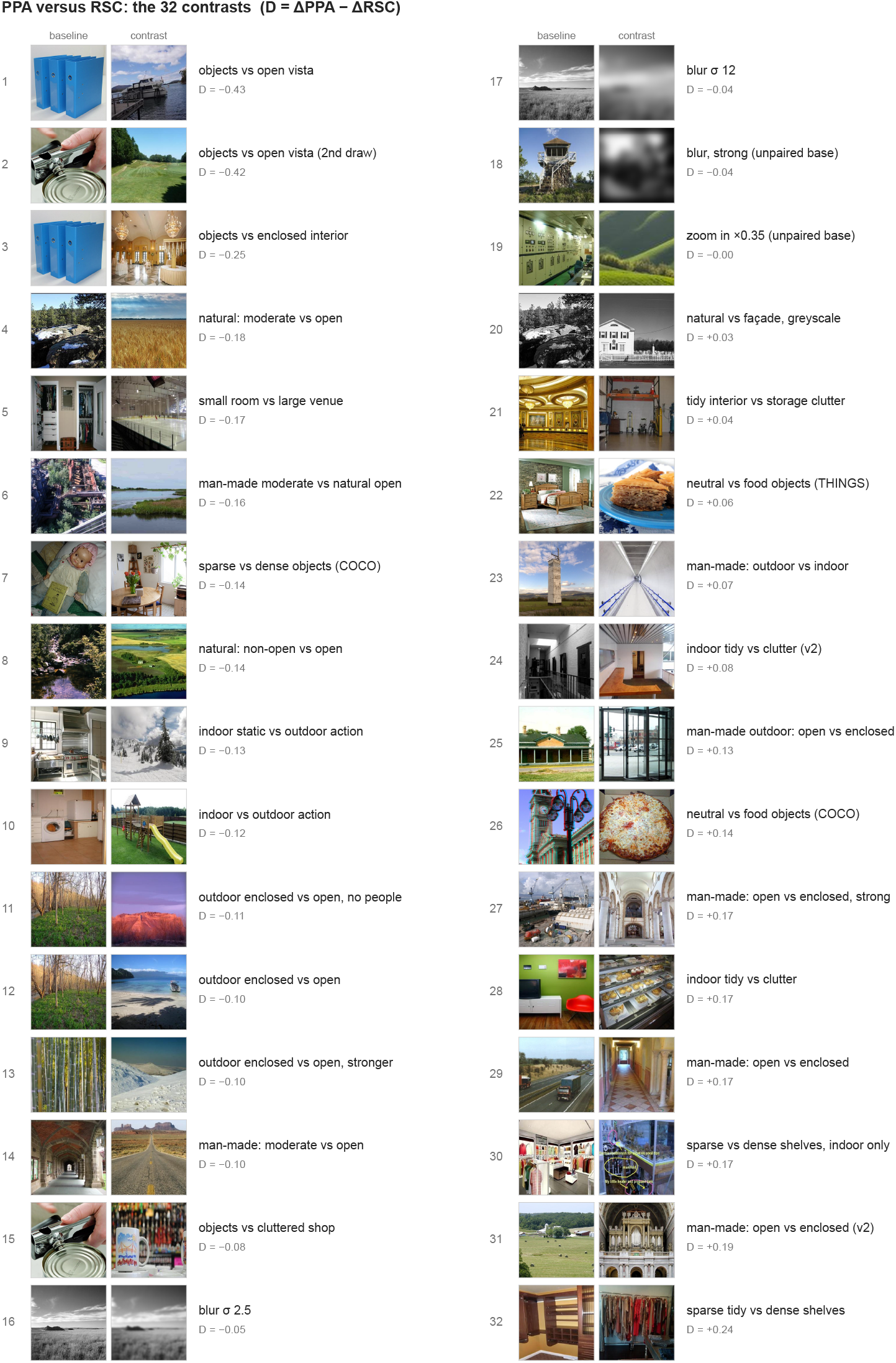
Braintree discovers stimulus manipulations that dissociate brain regions PPA and RSC.

We find it encouraging that the Braintree experiments naturally led to different kinds of experiments. Note also that many of the experiments Braintree tried were not successful and did not strongly separate the regions (Fig. 3a, c). Over successive experiments, Braintree identified several interpretable dimensions along which regions with otherwise similar category preferences respond differently. These results show that Braintree can adapt its experimental search to the scientific problem at hand, instead of simply executing a fixed set of experiments. In each case, it eventually identified some stimulus that produced different response in these region pairs, which gives us (another) concrete hypothesis to test experimentally in the human brain.

### 3.3 Braintree finds stimulus manipulations that dissociate ANN models

We next asked if Braintree could help with an open problem in NeuroAI. How do we tell models apart? We’re now at a stage where many candidate models of the brain predict brain responses about equally well (Schrimpf et al., 2020; Conwell et al., 2024). Can Braintree find interpretable stimulus dimensions for which these models make the most divergent, or *controversial*, predictions (inspired by: (Golan et al., 2020; 2026))? If successful, finding these stimulus dimensions would be diagnostic of how the models differ from each other and tell us which experiments would be informative to run when comparing models against the brain.

#### Braintree pipeline

For this experiment, Braintree was given a pair of brain models (CLIP-ResNet50 and CLIP-ViT-B/32) and asked to find stimulus manipulations that maximally dissociate their predicted responses. The orchestrator agent identifies candidate visual dimensions along which the models might differ. The planner agent turns each candidate into an experiment. The designer agent constructs controlled stimulus sets that vary the proposed dimension. After the stimuli are independently audited, the experimenter agent runs the experiment across the two models and measures how their predictions change. The resulting responses are passed through the fixed analysis pipeline and interpreted by the analyzer agent. Braintree then uses the observed model differences to refine what to test next, iteratively searching for stimulus dimensions that produce strong and interpretable model dissociations.

#### Results

Braintree was tasked with finding interpretable stimulus manipulations that produced different predictions from the two models. In total, Braintree tested 36 contrasts in PPA, 11 in FFA, and 15 in EBA. Braintree found multiple ways to separate the predictions of the two models (Appendix A.4). In PPA, for example, phase scrambling produced disagreements in both directions depending on its spatial scale. Scrambling with a coarse 8×8 grid of windows favored CLIP-ViT-B32 (D = −0.29 su, *p* = 1.7 × 10^−8^), while scrambling with a fine 32×32 grid favored CLIP-RN50 (D = 0.17 su, *p* = 4.7 × 10^−6^). In FFA, wild-animal faces strongly favored CLIP-ViT-B32 (D = −0.43 su, *p* = 4.6 × 10^−19^), and faces from FairFace favored CLIP-RN50 (D = 0.2 su, *p* = 1.9 × 10^−3^). In EBA, older men versus men produced a large disagreement toward CLIP-ViT-B32 (D = −0.47 su, *p* = 1.7 × 10^−11^), while children versus adults produced a similarly large disagreement toward CLIP-RN50 (D = 0.43 su, *p* = 1.3 × 10^−7^). We also observed that not every experiment was able to dissociate the model pair. Many manipulations proposed by Braintree changed the predictions of both models in similar ways. We implemented a stopping budget, which led to different regions having different numbers of experiments. Across all experiments, 28/62 manipulations produced reliable model dissociations (14/36 for PPA, 5/11 for FFA, 9/15 for EBA). The experiments that failed to dissociate the models were useful too. They ruled out plausible stimulus dimensions as major sources of disagreement and helped Braintree focus its search elsewhere.

Even though Braintree discovered these experiments by comparing only CLIP-RN50 and CLIPViT-B32, we next asked whether the discovered stimulus dimensions generalized beyond these two models. We therefore evaluated the same experiments on ten additional model backbones fit to each brain region. They did. Stimulus manipulations that produced larger disagreements between the original two models also produced greater variance across the broader set of models (Spearman *ρ* = 0.54, *n* = 62 contrasts, *p* = 6.6 × 10^−6^). Thus, experiments discovered from a single pair of models generalized to a much larger model population: the more strongly an experiment pulled the original models apart, the more it exposed differences among models more broadly. A detailed preview of the stimuli is provided in Appendix A.4, Fig. 7 and Fig. 8.

**Figure 7.**
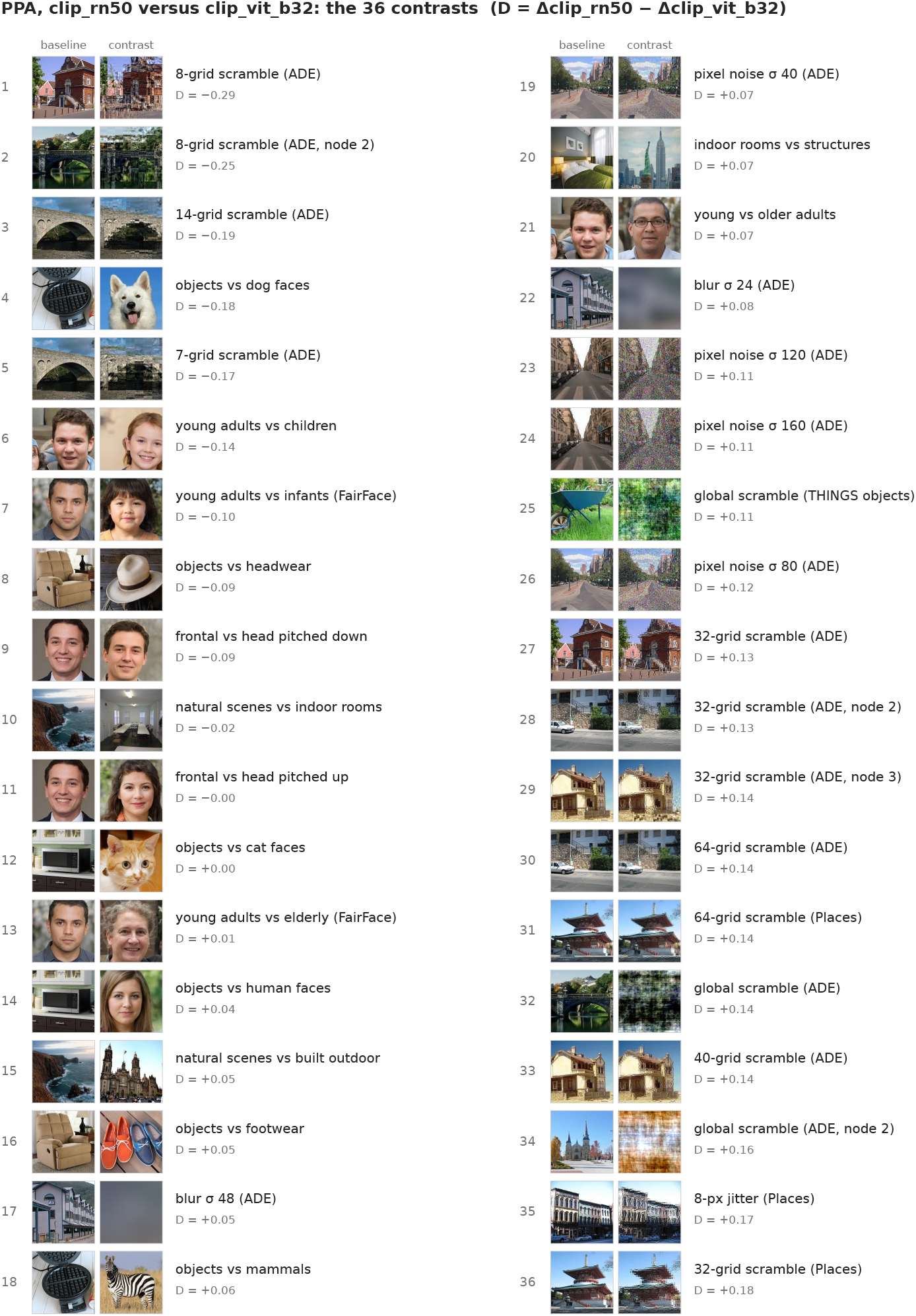
Braintree discovers stimulus manipulations that dissociate PPA models CLIP RN50 and CLIP VIT_b32. Human faces shown in this figure are synthetically generated.

**Figure 8.**
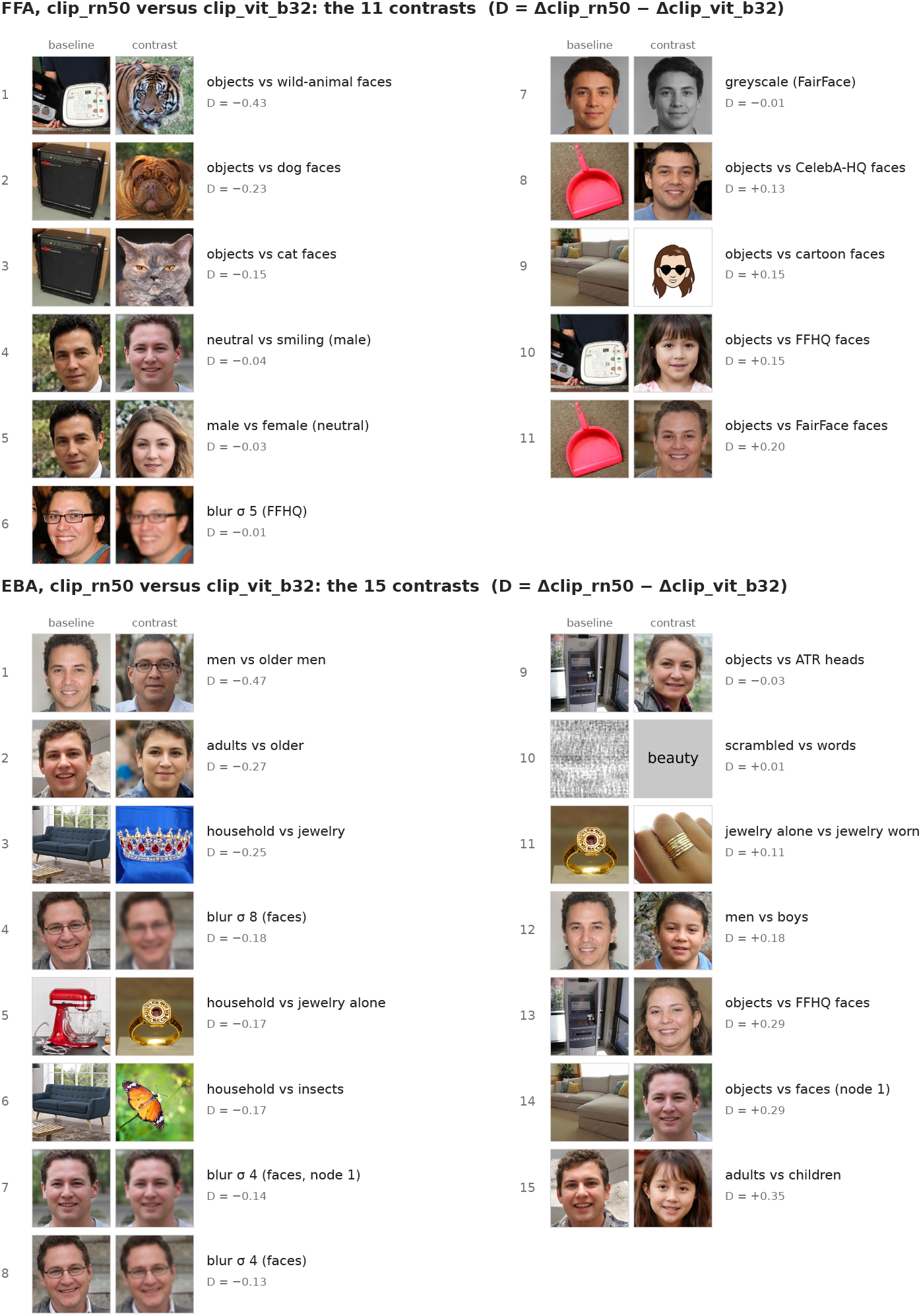
Braintree discovers stimulus manipulations that dissociate FFA models CLIP RN50 and CLIP VIT_b32, and dissociate EBA models CLIP RN50 and CLIP VIT_b32. Human faces shown in this figure are synthetically generated.

These results show how Braintree can help us understand where brain models differ. It can search for experiments that expose disagreements between models, including differences that may be hard to see from their overall prediction scores. Some models separate readily, while others make similar predictions across many experiments. The experiments Braintree finds give us concrete hypotheses to test in the brain and a way to ask which model better captures the neural response.

## 4 Discussion

We introduced Braintree, an agentic framework for running experiments in vision neuroscience. Braintree takes a scientific question through experimental design, stimulus construction, execution on in-silico models of the brain, analysis, and inference, while allowing the scientist to intervene at any stage. We showed that Braintree can reproduce published experiments (Section 3.1), replicate findings with new stimuli (Section 3.1), discover interpretable stimulus dimensions that dissociate brain regions (Section 3.2), and design experiments that expose disagreements between competing brain models (Section 3.3). We think these results showcase a fundamentally different way of doing vision science, where AI agents work together with stimulus-generation methods and in-silico models of the brain to design, run, and learn from experiments.

But there are limitations that are important to highlight. First, Braintree currently runs experiments on in-silico brain models. A dissociation found by the model is at this stage is only a *prediction* about the brain. We should consider them to be a (promising) testable hypothesis and confirm them on human neural measurements. Establishing how often these hypothesis generated from in-silico studies transfer to actual experiments is an important next step. Second, even though we have access to whole-brain models, our demonstrations in the paper focused on a small set of well-studied category-selective regions. There are many other brain regions and ideas to explore. Extending Braintree to whole-brain responses, dynamic stimuli, behavior, and/or to other species and measurements (like EEG) will require some additional experimental and analysis tools. But the Braintree design framework is modular, and new models, stimulus-generation methods, and measurement spaces can be easily added. Finally, agentic systems can make mistakes. Braintree separates experimental design, stimulus construction, quantitative analysis, and inference, and uses independent auditing agents. These safeguards reduce obvious failure modes, but they’re not perfect. Human inspection will be important, particularly as Braintree is applied to less established scientific questions.

Braintree is still a proof of principle, but we think it marks a step-change in how vision neuroscience *can* be done. Vision neuroscience is becoming *experiment-computable*, and we’ve shown how agents can now operate across much of the experiment loop. These ingredients make possible a new kind of in-silico neuroscience, where large parts of an experiment can be designed, executed, and analyzed directly on models. Braintree can remove many of the bottlenecks that limit what vision scientists can try, leaving room for human ideas, judgment, and intervention. If successful, systems like Braintree could let researchers explore a much larger space of experiments, identify the ones that are most informative, and then bring those experiments back to the brain. We think Braintree is the first step toward that future for vision neuroscience.

### AI use statement

Braintree is an agentic framework where language models participate in all stages of the scientific workflow. This includes reasoning over scientific questions, experimental design, stimulus construction, analysis, and interpretation. Evaluating the outputs of these agents is a central subject of this work. We therefore report agent-generated experimental designs, stimuli, analyses, and hypotheses as experimental outputs of the system.

Please note that all scientific claims and quantitative results reported in the paper were reviewed and verified by the *all* authors. We made sure that the quantitative outcomes used to evaluate experiments were computed using fixed analysis code (not by language models). Agent-generated experimental designs, stimuli, and interpretations were inspected by the authors, and the authors take full responsibility for the conclusions drawn from them. Generative AI (Claude) tools were used during development to assist with code, debugging, and refinement of experimental procedures. During manuscript preparation, generative AI was used only lightly to improve the clarity, grammar, and organization of our text.

We take full responsibility for the content in this work. This includes text, code, experimental designs, stimuli, claims, and other artifacts produced with the assistance of generative AI.

### Reproducibility statement

We have taken several steps to make Braintree and the experiments reported in this paper reproducible. The Appendix provides implementation details for the agentic pipeline, including the agents, models, and experimental protocols used in our experiments. We additionally provide details of the in-silico brain models, stimulus construction procedures, datasets, and quantitative analyses used for each experiment in Appendices A.1–A.4. All quantitative results are computed using fixed analysis code rather than language-model judgments. We will release the Braintree code, experimental configurations, generated stimuli, and analysis code needed to reproduce the experiments reported in this paper.

## Acknowledgments

This work was supported by a CoCo fellowship by the School of Psychology, Georgia Tech, to A.D.; the NIH Pathway to Independence Award (R00EY032603) from the National Eye Institute (NIH) and NSF Nexus (Allocation number: SOC250049) to N.A.R.M.

## A Appendix

### A.1 Braintree: Agentic Experimental Pipeline

Braintree is a multi-agent system for designing and conducting vision neuroscience experiments in silico. Given a scientific question or a published finding, Braintree can identify and ground the relevant hypothesis, design an experiment and construct its stimuli, run the experiment on image-computable models of the human visual system, and analyze and interpret the resulting responses. The resulting inference can then guide subsequent experiments. A user can inspect or intervene at any stage.

A central design principle of Braintree is to separate the different stages of the experimental process. Hypothesis formulation, experimental design, stimulus construction, execution, quantitative analysis, and scientific inference are handled by distinct agents or procedures, allowing the output of each stage to be independently checked before it is passed to the next. Auditor agents verify experimental designs, stimuli, analyses, and inferences throughout the pipeline. Quantitative outcomes are additionally isolated from language-model reasoning. All statistics are computed from the predicted neural responses using fixed analysis code. This modular structure makes each stage of an experiment independently inspectable and reproducible.

Figure 1 provides an overview of the system.

#### A.1.1 Agent architecture

Braintree uses an orchestrator together with a set of specialized agents. The orchestrator maintains the scientific goal of the study and determines which agent should act next. Specialized agents are responsible for different parts of the experimental process: the scientific reasoning agent understands and grounds the scientific question; the experiment planner and stimulus designer construct the experiment and its stimuli; the experimentalist executes the experiment; and the analyst and inference agent analyze and interpret the resulting responses. Independent auditor agents operate across these stages.

We deliberately separate agents that *produce* an experimental object from agents that *evaluate* it. For example, the stimulus designer that constructs a stimulus set does not certify its quality. A separate auditor independently checks whether the stimuli implement the intended experimental manipulation. Similarly, the inference agent that interprets experimental results does not compute the statistical outcome. All quantitative outcomes are instead computed using fixed analysis code.

The agents communicate through structured experiment records rather than relying on a single shared conversation. Experimental specifications, stimulus sets, analyses, and decisions are saved between stages. This allows individual agents to operate with limited context and provides a record of how each experiment was produced.

Agents communicate through structured artifacts that record the output of each stage of an experiment. Let *q* denote the research question, *g* the grounded experimental goal, *E* the executable experimental specification, *D* the resulting neural response data, and *f* the scientific inference. The flow of information is therefore

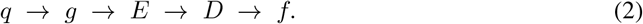

Each artifact is persisted and can be independently inspected or audited before being passed to subsequent stages.

#### A.1.2. Experimental safeguards

Allowing agents to design experiments creates an obvious risk: the same system could design an experiment, inspect its outcome, and then change the experiment until it obtains a desired result. Braintree uses several safeguards to reduce this flexibility.

##### Separate production and evaluation

Agents that construct stimuli or experiments do not evaluate their own outputs. Independent auditing agents check stimulus fidelity and experimental specifications before execution.

##### Fixed quantitative analyses

Statistical outcomes are computed by deterministic analysis code rather than by language models. The agents receive these computed results and can interpret them, but cannot alter the underlying measurements or statistical decision.

##### Experiments are specified before execution

The experimental conditions, primary comparisons, and predicted outcomes are recorded before the corresponding neural responses are computed. Changes to an experiment produce a new experimental specification rather than silently modifying the previous one.

##### Stimuli are not selected using their predicted responses

Images used for confirmatory experiments are selected or generated without observing their predicted neural responses. When model responses are explicitly used to search for informative stimuli, discovery and evaluation are performed using separate model configurations or held-out stimulus sets.

##### Complete provenance

Braintree records the experimental specification, stimuli, model configuration, analysis, and agent outputs associated with each experiment. This allows an experimental result to be traced back to the exact inputs and models that produced it.

These safeguards are implemented in code where possible. For example, the execution pipeline can reject an experiment whose stimuli have not passed the required audit. We use language-model instructions primarily for scientific reasoning and reserve deterministic checks for constraints that directly affect experimental validity.

### A.2 In-silico models of the brain

Braintree conducts experiments on in silico image-computable encoding models of the human visual cortex. Each model takes an image as input and predicts fMRI responses for all voxels in the brain. We trained models independently for four participants (subjects 1, 2, 5, and 7) from the Natural Scenes Dataset (NSD), using responses to the 1,000 COCO images seen by all participants. We used the 1.8 mmş native surface preparation and version 3 single-trial beta estimates (betas_fithrf_GLMdenoise_RR). To reduce session-to-session variability, voxel responses were zscored within each session and then averaged across the three repetitions of each stimulus. We focus on category-selective regions of interest (ROIs): FFA (FFA-1 and FFA-2), PPA, EBA, OFA, and RSC. Each ROI is defined per participant from the NSD category localizer, keeping voxels with *t >* 7, and a model’s ROI response is the mean over those voxels.

Our primary encoding models use one of two visual backbones: DINOv2 ViT-L/14, a selfsupervised vision transformer, or the ResNet-50 image encoder of CLIP, a convolutional network trained with language supervision. Each is followed by a linear readout that takes features from a single intermediate layer, chosen per ROI by a layer sweep on validation data. The readout’s first step depends on the backbone’s feature format: for DINOv2, projections over feature channels and over tokens form a 2,048-dimensional bottleneck; for CLIP-RN50, a 3 × 3 convolution reduces the feature map to 128 channels, which are averaged over space. In both cases, the bottleneck is mapped linearly to voxels, with no nonlinearity, and trained with Adam on mean squared error. Images are square-padded, resized to 224 ×224, and normalized with each backbone’s own statistics (ImageNet for DINOv2, OpenAI CLIP for CLIP-RN50).

The encoding models used in this work (CLIP RN50, CLIP ViT-B/32, DINOv2, BLIP-2, CLIP RN101, Kosmos-2, EfficientNet, DINOv2-L, SigLIP 2, BRIA, EVA-02, HRNet, and FBNet) achieve mean univariate Pearson accuracies of 0.704 (variance 0.014) across five brain regions (EBA, FFA, OFA, PPA, and RSC). Within each ROI the thirteen models are tightly clustered, with standard deviations of 0.012, 0.011, 0.015, 0.006, and 0.010.

### A.3 Reproducing and replicating published experiments

We evaluate whether Braintree can recover established findings from the vision neuroscience literature. Given a published paper, Braintree identifies a central claim, the brain region involved, the experimental comparison supporting the claim, and the principal control used to distinguish the proposed explanation from alternatives.

Braintree then reconstructs the corresponding experiment. We distinguish two settings. In a *reproduction*, Braintree uses the original stimuli released by the authors whenever they are available. In a *replication*, Braintree constructs a new stimulus set implementing the same experimental comparison. This distinction tests separately whether the in-silico models recover the reported result on the original experiment and whether the effect generalizes to newly constructed stimuli.

For each experiment, the planner specifies the experimental conditions and primary contrast before model responses are measured. The stimulus designer then constructs the required image sets using existing annotated datasets, algorithmic image manipulations, or generative image methods as appropriate. An independent auditor checks whether the resulting images implement the requested manipulation and whether unintended differences between conditions are apparent.

Once the stimuli pass this check, they are presented to the encoding models. Predicted responses are analyzed using fixed statistical routines, and the resulting measurements are returned to the inference agent for scientific interpretation.

Table 3 lists the published findings used to evaluate Braintree’s ability to reproduce prior vision neuroscience experiments. The studies span three category-selective regions (PPA, FFA, and EBA) and test a range of established response properties.

Table 4 lists the published findings used to evaluate Braintree’s ability to replicate prior vision neuroscience experiments. The studies span three category-selective regions (PPA, FFA, and EBA) and test a range of established response properties.

### A.4 Quantifying the dissociation between ROI pairs and model pairs

To compare readouts with different response scales, we express predictions in standard units (su). For a single readout (one model, one participant, one ROI), let *p*(*i*) be the predicted ROI response to image *i*. We rescale it by the readout’s own mean responses to two fixed reference sets of 20 images each: a preferred-category set P and a non-preferred set N (FFA: faces and places; PPA: places and faces; EBA: bodies and places):

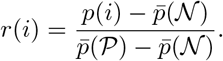

A value of 0 is the readout’s mean response to the non-preferred category, and 1 is its mean response to the preferred category, so one su equals that readout’s own category selectivity.

Each manipulation compares a base image set *B* with a changed set *E*. The change in readout *x* is 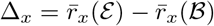, and the dissociation between readouts *A* and *B* is

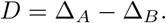

A positive *D* means the manipulation drives *A* more than *B*. We compute *D* within each participant and average it across the four participants.

For statistical tests, images are the unit of observation. We first average each image’s responses across participants; since *D* is a difference of means, this leaves its value unchanged. The test depends on how the two image sets are related. If *E* and *B* contain different images, we score each image by how much more it drives *A* than *B, u*_*i*_ = *r*_*A*_(*i*) − *r*_*B*_(*i*); then *D* = ū(E) − *ū*(B), and we compare the two sets with a Welch two-sample *t*-test. If E contains edited versions of the images in B, we score each baseedit pair by how much more the edit changes *A* than *B, d*_*i*_ = [*r*_*A*_(*e*_*i*_) − *r*_*A*_(*b*_*i*_)] − [*r*_*B*_(*e*_*i*_) − *r*_*B*_(*b*_*i*_)]; then 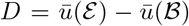, and we use a one-sample *t*-test. Because the encoding models are deterministic, almost any nonzero difference is significant against zero. We therefore test against a smallest effect of interest, *δ* = 0.10 su (10% of the readout’s category selectivity), using one-sided tests: *D > δ* with *t* = (*D* − *δ*)*/*SE to conclude that the manipulation drives *A* more than *B*, and *D <* −*δ* with *t* = (−*δ* − *D*)*/*SE to conclude the reverse.

To ask whether a dissociation between *A* and *B* reflects a broader disagreement among models, we also measure each manipulation on a set of ten backbones: BLIP-2, CLIP-RN101, Kosmos-2, EfficientNet, DINOv2-L, SigLIP2, BRIA, EVA-02, HRNet, and FBNet. Each backbone has its own readouts, so its change Δ_*m*_ is expressed in its own standard units. For each manipulation, we measure the divergence among models as the variance of their changes, Var_*m*_(Δ_*m*_), across the ten backbones. We then relate this divergence to |*D*| with a Spearman correlation across all contrasts and ROIs.

